# *In vitro* Evaluation of Various Antimicrobials against Field *Mycoplasma gallisepticum* and *Mycoplasma synoviae* Isolates in Egypt

**DOI:** 10.1101/726000

**Authors:** Marwa I Abd El-Hamid, Naglaa FS Awad, Usama H. Abo-Shama, MH Yousreya, Mahmoud A Abdel-Rahman, Helal F. Hetta, Adel M Abdelaziz

## Abstract

Among many avian mycoplasmas, *Mycoplasma gallisepticum* (MG) and *Mycoplasma synoviae* (MS) are recognized as the main etiological agents of respiratory diseases and infectious synovitis in chickens and turkeys causing tremendous economic losses worldwide. Therefore, proper treatment is promoted for the control of these diseases. This study was the first in Egypt to evaluate the *in vitro* efficacy of various antimicrobials against field MG and MS isolates recovered from chicken and turkey flocks using both conventional microdilution and quantitative real-time PCR (qRT-PCR) assays. Totally, 47 mycoplasma isolates were recovered from 160 collected tracheal samples (29.4%). Of these, 44 MG (27.5%) and 3 MS (1.9%) were identified using conventional and molecular assays. The *in vitro* susceptibilities of 4 representative mycoplasma isolates (3 MG and one MS) to 8 antibiotics and 4 essential oils were investigated. The tested isolates showed various susceptibilities to tested antimicrobials. Toldin CRD, followed by clove, cumin and cinnamon oils were commonly effective against both MG and MS clinical isolates with MIC values ranging from 0.49 to 15.63 µg/mL. Similarly, tylvalosin was the most active antibiotic against MG and MS isolates with the lowest MIC values (0.015-0.03 µg/mL). DNA copies of both MG *mgc2* and MS *vlhA* genes were markedly decreased upon treatment with majority of tested antimicrobials confirming their effectiveness as was also evaluated by conventional MIC results. In conclusion, Toldin CRD and tylvalosin were found to be the most effective antimicrobials in this study, which will contribute in controlling avian mycoplasma infections.

**Author Summary:** Avian mycoplasmosis is considered as one of the most prominent economic problems in the commercial poultry industry worldwide. Antimicrobial therapy is the most effective tool for treatment of mycoplasmas. Owing to the side effects of antibiotics and the development of resistance to the currently used drugs, an increased emphasis on the use of alternative antimicrobials is of utmost importance. Here, we evaluate the *in vitro* inhibitory effects of some essential oils and various commercial antibiotics against *Mycoplasma gallisepticum* (MG) and *Mycoplasma synoviae* (MS) field isolates using micro-broth dilution method and qRT-PCR assays. We found that toldin CRD, followed by clove, cumin and cinnamon oils were effective against both MG and MS clinical isolates. Similarly, tylvalosin was the most active antibiotic against MG and MS isolates. We also found that DNA copies of both MG *mgc2* and MS *vlhA* genes were markedly decreased upon treatment with majority of tested antimicrobials. Our study provides new insights into the control of avian mycoplasma infections.

## Introduction

Avian mycoplasmosis has been deemed as one of the most prominent economic problems in the commercial poultry industry all over the world. More than 20 species of genus *Mycoplasma* are known to infect avian hosts with *Mycoplasma gallisepticum* (MG) and *Mycoplasma synoviae* (MS) being the most clinically relevant mycoplasmas [1].

*Mycoplasma* species are transmitted horizontally by susceptible birds through direct or indirect contact with contaminated surfaces [2] and vertically from dam to offspring through eggs [3].

In geographic localities where mycoplasmosis is enzootic, programs for controlling their infections are the most practical ways to decrease the resulted economic losses. Antimicrobial therapy remains the most prompt and effective tool for treatment of mycoplasmas in poultry farms. Basically, MG and MS have shown *in vitro* and *in vivo* susceptibilities to several antimicrobials such as macrolides, tetracyclines and quinolones [4,5], but they are resistant to penicillins or other antibiotic inhibitors of cell wall synthesis because of their cell-wall deficiency.

Veterinarians usually ignore the side effects of synthetic antibiotics while recommending the therapy for avian mycoplasmosis. Moreover, the development of resistance to currently used drugs [1,6] incorporating with their high costs increased emphasis on the use of new and alternative antimicrobials.

An approach for the development of new antimicrobials mainly from plant origin offers a great contribution in the fight against mycoplasma infections. To our knowledge, few researchers reported the efficacy of medicinal plants against different species of mycoplasma [7,8].

Performance of phenotypic antimicrobial susceptibility tests for mycoplasmas is time-consuming and requires special techniques. Therefore, quantitative real time PCR (qRT-PCR) assay has been recently developed as a faster, easier, sensitive and more specific method compared with conventional ones used in clinical microbiology laboratories [9].

Based on the above background, this study was conducted to evaluate the *in vitro* inhibitory effects of some essential oils and various commercial antibiotics against MG and MS field isolates using micro-broth dilution method and qRT-PCR assays.

## Materials and methods

### Sample collection and study area

The current study was carried out from January 2017 to December 2018. Three commercial broiler chicken flocks located in Faiyum, Giza and Sharkia Governorates, Egypt as well as one turkey flock in El Dakhleya Governorate, Egypt were examined. The birds from Giza (50), Sharkia (50) and El Dakhleya (10) were suffering from respiratory manifestations in the form of conjunctivitis, nasal and ocular discharges, sneezing, coughing, rales, gasping, depression and weakness, while those from Faiyum were suffering from both respiratory disease and synovitis. All of these flocks were not vaccinated against any mycoplasma diseases. From each flock, tracheal swabs from all diseased birds were aseptically collected and transferred in test tubes with Frey’s broth medium as a transport medium to the laboratory until being subjected to MG and MS isolation and identification.

### Ethical statement

All animal protocols were approved by the Institutional Animal Care and Use Committee (IACUC), Faculty of Veterinary Medicine, Zagazig University.

### Isolation and identification of *M. gallisepticum* and *M. synoviae*

After incubating the broth media for 3-5 days, the broth culture was streaked onto Frey’s agar medium and incubated for 21 days. Inoculated broth and agar media were incubated under microaerophilic humid conditions (10% CO_2_) at 37ºC. When the growth of the colonies was obtained, digitonin test was performed. Identification of MG and MS was made by a combination of conventional biochemical methods (glucose fermentation, arginine deamination and film and spot formation tests) and serologically via growth inhibition test using specific antisera [10]. Further molecular identification of both MG and MS was carried out as was described previously by PCR amplifications of *mgc2* and *vlhA* genes, respectively [11,12]. Known positive MG and MS reference strains as well as negative controls were included in every set of PCR reactions. The sizes of expected amplification products were 236-302 bp for *mgc2* and 1.1 kbp for *vlhA* gene.

### *Mycoplasma* reference strains

*Mycoplasma gallisepticum* and *Mycoplasma synoviae* reference strains with accession numbers of HQ591357 and KT943466 were used for all PCR assays.

### Natural substances

The essential oils of clove (*Syzygium aromaticum*), cinnamon (*Cinnamomum zeylanicum*) and cumin (*Cuminum cyminum*) were purchased from National Research Centre, Egypt. Additionally, Toldin CRD (B.NO, 00101) which is a prophylactic and therapeutic agent for chronic respiratory disease (CRD) complex in poultry was kindly supplied by Dawa International for Pharmaceutical Industries, Egypt. It contains many active natural compounds such as aliumcepa, ginger, cinnamon, oregano, circumin, liqurice, anise oils and propolis.

### Antibiotics

The following 8 antibiotic agents of 5 different groups; macrolides, fluoroquinolones, tetracyclines, lincosamide and pleuromutilins were used for the susceptibility tests: tylvalosin (Pharmgate Animal Health Canada, Inc), tilmicosin (ELANCO, Geneva), tylosin (ELANCO, USA), enrofloxacin (INVESA, Spain), doxycycline (Oxoid, UK), chlortetracycline (Oxoid, UK), lincomycin (Oxoid, UK) and tiamulin (Sandoz GmbH, Basel, Switzerland).

### Antimicrobial susceptibility testing of mycoplasma isolates using microdilution and qRT-PCR assays

The antimicrobial susceptibility rates of four representative mycoplasma field isolates (one isolate from each flock) were determined by both conventional broth microdilution [13] and qRT-PCR [14] assays.

Briefly, each tested antimicrobial substance was twofold serially diluted starting from a concentration of 1024 µg /mL for each antibiotic and 125 µg/mL for each essential oil. Mycoplasma isolates were also diluted to contain 10^3^-10^4^ color changing unit/0.2 mL. Consequently, 0.1 mL volume of each diluted substance was mixed with 0.1 mL of each diluted mycoplasma isolate in the 96-well microtiter plates. Growth controls (tested field mycoplasma strains grown into broth medium without any tested substances), sterility controls (broth medium without neither tested substances nor mycoplasma inoculum) and pH control (broth medium adjusted to pH 6.8) were included in each plate. The plates were sealed, incubated at 37° C and inspected at regular intervals for any changes in the color indicator. Each experiment was performed in triplicate and repeated twice to confirm the results. In accordance with the previously mentioned guidelines [13], the initial MIC was read when the change of color in the broth was first observed in the growth control wells. The final MIC was read at the time when no further color change in the wells containing antimicrobials was observed.

At 1^st^, 2^nd^, 3^rd^ and 4^th^ weeks after the incubation, 50 µL aliquots from each well was harvested and subjected for determination of DNA loads by qRT-PCR using MG and MS specific primers. The number of DNA copies at all intervals was determined using standard curves prepared from tenfold serial dilution of genome copies of MG and MS reference strains. After amplification, a final melting curve analysis was performed to determine the presence or absence of non-specific amplification products. The inhibition rates of all tested substances were calculated according to the following formula: Inhibition rate (%) = [(average of DNA loads in control wells-DNA load in test well) / average of DNA loads in control wells] x 100 [14]. The MIC of each tested substance was defined as the lowest concentration causing 99% inhibition of mycoplasma field isolates.

Interpretation of antimicrobial susceptibility results is hampered by lacking of the official breakpoints. In these cases, interpreting the data of our study is based on values previously used in other publications [8,15].

## Results

### Prevalence and characterization of avian *M. gallisepticum* and *M. synoviae*

Fried-egg-shaped pure colonies of cultured mycoplasmas were seen on Frey’s agar medium. All the colonies were found sensitive to 1.5% digitonin, ensuring that they were mycoplasmas.

The recovery rates of avian *Mycoplasma* species from 4 flocks located in different localities are illustrated in Table 1. A total of 47 mycoplasma isolates out of 160 collected tracheal samples (29.4%) were identified biochemically into 2 groups. The first group was positive for glucose fermentation test and negative for both arginine deamination and film and spot formation tests, while the second group fermented glucose, did not hydrolyse arginine and was positive for the film and spot formation test. According to serological identification results using growth inhibition test, 44 (27.5%) and 3 (1.9%) were inhibited by specific antisera to MG and MS, respectively. Moreover, PCR confirmed the nucleic acid of all biochemically and serologically identified isolates with the production of the predicted amplicons to MG and MS at 236-302 bpand 1.1 kbp, respectively. The total recovery rate of *Mycoplasma* species from the examined chicken and turkey flocks were 29.3% (44/150) and 30% (3/10), respectively. Cumulatively, phenotypic and genotypic identification results confirmed that 41 isolates (27.3%) of 150 tracheal swabs from chicken flocks and 3 isolates (30%) of 10 tracheal swabs from the turkey flock in El Dakhleyawere MG. In addition, MS were recovered from the chicken flocks with a percentage of 2% (3/150).

**Table 1.**
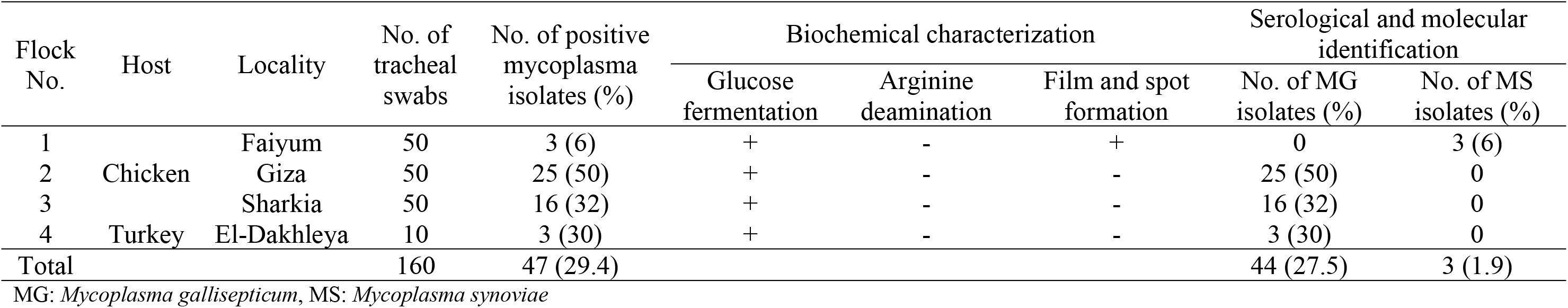
Prevalence and characterization of avian *M. gallisepticum* and *M. synoviae* in 4 flocks from different localities

| Flock No. | Host | Locality | No. of tracheal swabs | No. of positive mycoplasma isolates (%) | Biochemical characterization |  |  | Serological and molecular identification |  |
| --- | --- | --- | --- | --- | --- | --- | --- | --- | --- |
|  |  |  |  |  | Glucose fermentation | Arginine deamination | Film and spot formation | No. of MG isolates (%) | No. of MS isolates (%) |
| 1 | Chicken | Faiyum | 50 | 3 (6) | + | - | + | 0 | 3 (6) |
| 2 |  | Giza | 50 | 25 (50) | + | - | - | 25 (50) | 0 |
| 3 |  | Sharkia | 50 | 16 (32) | + | - | - | 16 (32) | 0 |
| 4 | Turkey | El-Dakhleya | 10 | 3 (30) | + | - | - | 3 (30) | 0 |
| Total |  |  | 160 | 47 (29.4) |  |  |  | 44 (27.5) | 3 (1.9) |

MG: *Mycoplasma gallisepticum*, MS: *Mycoplasma synoviae*

### *In vitro* antimicrobial activities of essential oils and antibiotics

#### Conventional broth microdilution assay

In the first part of the study, 4 selected mycoplasma isolates (3 MG and one MS) representative for 4 flocks located in different localities were investigated. The *in vitro* activities of 4 different essential oils and 8 antibiotics of 5 groups against these isolates were detected by the microbroth dilution technique as illustrated in Tables 2–4. Herein, MIC initial values were those used to evaluate the susceptibility of examined mycoplasma isolates to different antimicrobials. The *in vitro* MIC results of selected natural products revealed that clove, cinnamon, cumin and Toldin CRD essential oils were variously capable for inhibiting the growth of both MG and MS clinical isolates. Toldin CRD was found to be the most effective natural substance against all tested isolates with MIC values of 0.49-0.98 µg/mL. Moreover, both clove and cumin oils had excellent antimycoplasmal activities with MIC ranges of 0.49-1.95µg/mL and 0.98-3.91 µg/mL, respectively. On the other hand, cinnamon oil showed various activities from excellent to good against mycoplasma isolates with MIC values of 1.95-15.63 µg/mL with only one MG isolate showing an MIC value of 15.63 µg/mL.

**Table 2.**
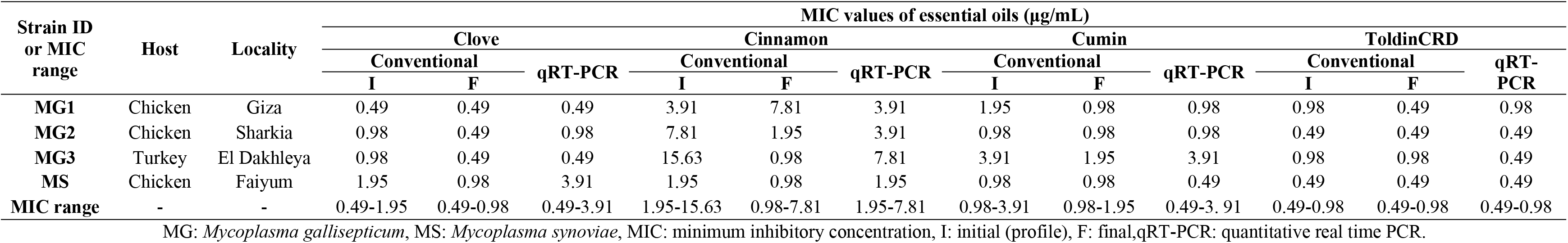
*In vitro* antimicrobial activities of four essential oils against *M. gallisepticum* and *M. synoviae* isolates as determined by conventional broth microdilution and qRT-PCR methods

| Strain ID<br>or MIC<br>range | Host | Locality | MIC values of essential oils (µg/mL) |  |  |  |  |  |  |  |  |  |  |  |
| --- | --- | --- | --- | --- | --- | --- | --- | --- | --- | --- | --- | --- | --- | --- |
|  |  |  | Clove |  |  | Cinnamon |  |  | Cumin |  |  | ToldinCRD |  |  |
|  |  |  | Conventional |  | qRT-PCR | Conventional |  | qRT-PCR | Conventional |  | qRT-PCR | Conventional |  | qRT-PCR |
|  |  |  | I | F |  | I | F |  | I | F |  | I | F |  |
| MG1 | Chicken | Giza | 0.49 | 0.49 | 0.49 | 3.91 | 7.81 | 3.91 | 1.95 | 0.98 | 0.98 | 0.98 | 0.49 | 0.98 |
| MG2 | Chicken | Sharkia | 0.98 | 0.49 | 0.98 | 7.81 | 1.95 | 3.91 | 0.98 | 0.98 | 0.98 | 0.49 | 0.49 | 0.49 |
| MG3 | Turkey | El Dakhleya | 0.98 | 0.49 | 0.49 | 15.63 | 0.98 | 7.81 | 3.91 | 1.95 | 3.91 | 0.98 | 0.98 | 0.49 |
| MS | Chicken | Faiyum | 1.95 | 0.98 | 3.91 | 1.95 | 0.98 | 1.95 | 0.98 | 0.98 | 0.49 | 0.49 | 0.49 | 0.49 |
| MIC range | - | - | 0.49-1.95 | 0.49-0.98 | 0.49-3.91 | 1.95-15.63 | 0.98-7.81 | 1.95-7.81 | 0.98-3.91 | 0.98-1.95 | 0.49-3. 91 | 0.49-0.98 | 0.49-0.98 | 0.49-0.98 |

MG: *Mycoplasma gallisepticum*, MS: *Mycoplasma synoviae*, MIC: minimum inhibitory concentration, I: initial (profile), F: final, qRT-PCR: quantitative real time PCR.

**Table 3.** Summary of initial and final MIC ranges (μg/mL) of the isolated *M. gallisepticum* and *M. synoviae* isolates with the suggested non-official breakpoints

| Antibiotic class | Antibiotic agent | Non-official<br>breakpoints<br>[15] | Initial MIC<br>range | Final MIC<br>range |
| --- | --- | --- | --- | --- |
| **Macrolides** | Tylvalosin | $S \leq 0.5$; $R > 2$ | 0.015-0.03 | 0.015-0.125 |
| | Tilmicosin | $S \leq 8$; $R \geq 32$ | 0.06-0.25 | 0.25-0.50 |
| | Tylosin | $S \leq 1$; $R \geq 4$ | 0.06 | 0.06-0.25 |
| **Fluoroquinolones** | Enrofloxacin | $S \leq 0.5$; $R \geq 2$ | 0.25-2 | 1-2 |
| **Tetracyclines** | Doxycycline | $S \leq 4$; $R \geq 16$ | 0.125-0.5 | 0.25-1 |
| | Chlortetracycline | $S \leq 4$; $R \geq 16$ | 0.5-16 | 2-16 |
| **Lincosamide** | Lincomycin | $S \leq 2$; $R \geq 8$ | 0.25-2 | 2-8 |
| **Pleuromutilins** | Tiamulin | $S \leq 8$; $R \geq 16$ | 0.06-0.5 | 0.5 |

S: susceptible, R: resistant

**Table 4.** Comparison between MICs of various antibiotics for *M. gallisepticum* and *M. synoviae* isolates as determined by conventional broth microdilution and qRT-PCR methods with their corresponding susceptibility patterns

| Strain ID | MIC values (µg/mL) |  |  |  |  |  |  |  |  |  |  |  |  |  |  |  |  |  |  |  |  |  |  |  |
| --- | --- | --- | --- | --- | --- | --- | --- | --- | --- | --- | --- | --- | --- | --- | --- | --- | --- | --- | --- | --- | --- | --- | --- | --- |
|  | TVN |  |  | TIL |  |  | TYL |  |  | EFX |  |  | DX |  |  | CTC |  |  | LCM |  |  | TIA |  |  |
|  | Conventional |  | qRT-PCR | Conventional |  | qRT-PCR | Conventional |  | qRT-PCR | Conventional |  | qRT-PCR | Conventional |  | qRT-PCR | Conventional |  | qRT-PCR | Conventional |  | qRT-PCR | Conventional |  | qRT-PCR |
|  | I | F |  | I | F |  | I | F |  | I | F* |  | I | F |  | I | F |  | I | F* |  | I | F |  |
| MG1 | 0.015 (S) | 0.125 | 0.031 | 0.125 (S) | 0.25 | 0.5 | 0.06 (S) | 0.25 | 0.125 | 0.25 (S) | 1 | 0.5 | 0.125 (S) | 1 | 0.25 | 0.5 (S) | 4 | 4 | 0.25 (S) | 2 | 0.25 | 0.125 (S) | 0.5 | 0.25 |
| MG2 | 0.03 (S) | 0.125 | 0.062 | 0.06 (S) | 0.5 | 0.5 | 0.06 (S) | 0.25 | 0.25 | 0.5 (S) | 2 (R) | 0.5 | 0.5 (S) | 1 | 0.5 | 16 (R) | 16 | 16 | 2 (S) | 8 (R) | 1 | 0.06 (S) | 0.5 | 0.5 |
| MG3 | 0.015 (S) | 0.015 | 0.031 | 0.25 (S) | 0.25 | 0.25 | 0.06 (S) | 0.06 | 0.125 | 2 (R) | 2 | 1 | 0.125 (S) | 0.25 | 0.125 | 1 (S) | 2 | 1 | 1 (S) | 2 | 2 | 0.25 (S) | 0.5 | 0.125 |
| MS | 0.015 (S) | 0.015 | 0.062 | 0.25 (S) | 0.25 | 0.5 | 0.06 (S) | 0.06 | 0.06 | 0.5 (S) | 2 (R) | 0.5 | 0.25 (S) | 1 | 0.25 | 2 (S) | 4 | 4 | 2 (S) | 8 (R) | 2 | 0.5 (S) | 0.5 | 0.5 |

MG: *Mycoplasma gallisepticum*, MS: *Mycoplasma synoviae*, MIC: Minimum inhibitory concentration, TVN: Tylvalosin, TIL: Tilimicosin, TYL: Tylosin, EFX: Enrofloxacin, DX: Doxycycline, CTC: Chlortetracycline, LCM: Lincomycin, TIA: Tiamulin, I: initial (profile), F: final, qRT-PCR: quantitative real time PCR, \*: Final MIC profiles differing from the initial ones, S: susceptible, R: resistant

The broadest ranges of MIC values were detected for antibiotics of the macrolide group with MICs ranging from 0.015-0.25 µg/mL. Among the examined 3 macrolides (tylvalosin, tilmicosin and tylosin), tylvalosin was found to be the most active antibiotic in the examination with the lowest MIC values (0.015-0.03 µg/mL) against MG isolates and 0.015 µg/mL against the MS isolate. Moreover, tilmicosin and tylosin demonstrated also lower MICs against both MG and MS isolates (up to 0.06 µg/mL each). From the pleuromutilins, the MIC values of tiamulin were low (0.06-0.5µg/mL). Among tetracyclines, the MIC values of both doxycycline and chlortetracycline showed a wide range (0.125 to 0.5 µg/mL and 0.5 to 16 µg/mL, respectively).

It was obviously noted that MS isolate was sensitive to all tested antibiotics. On the other hand, all MG isolates were sensitive to tylvalosin, tilmicosin, tylosin, doxycycline, lincomycin and tiamulin antibiotics and only 33.3% of the isolates were sensitive to each of enrofloxacin and chlortetracycline.

A Comparison of initial and final MIC values showed zero to three-fold differences for all antibiotics. Only in the case of 6 antibiotics (tylvalosin, tilmicosin, doxycycline, chlortetracycline, lincomycin and tiamulin), initial and final MICs exhibited up to 3 fold differences among MG isolates. Values for initial and final MICs of tylvalosin, tilmicosin, tylosin and enrofloxacin and tylvalosin, tilmicosin, tylosin and tiamulin are the same for each of MG3 and MS isolates, respectively. Final MICs for enrofloxacin and lincomycin differed from those of the initial MICs among each of MG2 and MS isolates, including changes of MIC profiles from sensitive to resistant.

#### Quantitative real-time PCR assay

In the second part of our work, we quantitatively assessed the *mgc2* and *vlh1* gene copies when MG and MS isolates were grown in the presence of all tested antimicrobials at 1^st^, 2^nd^, 3^rd^ and 4^th^ weeks of incubation, respectively (Tables 2 and 4, Figure 1).

**Fig. 1.**
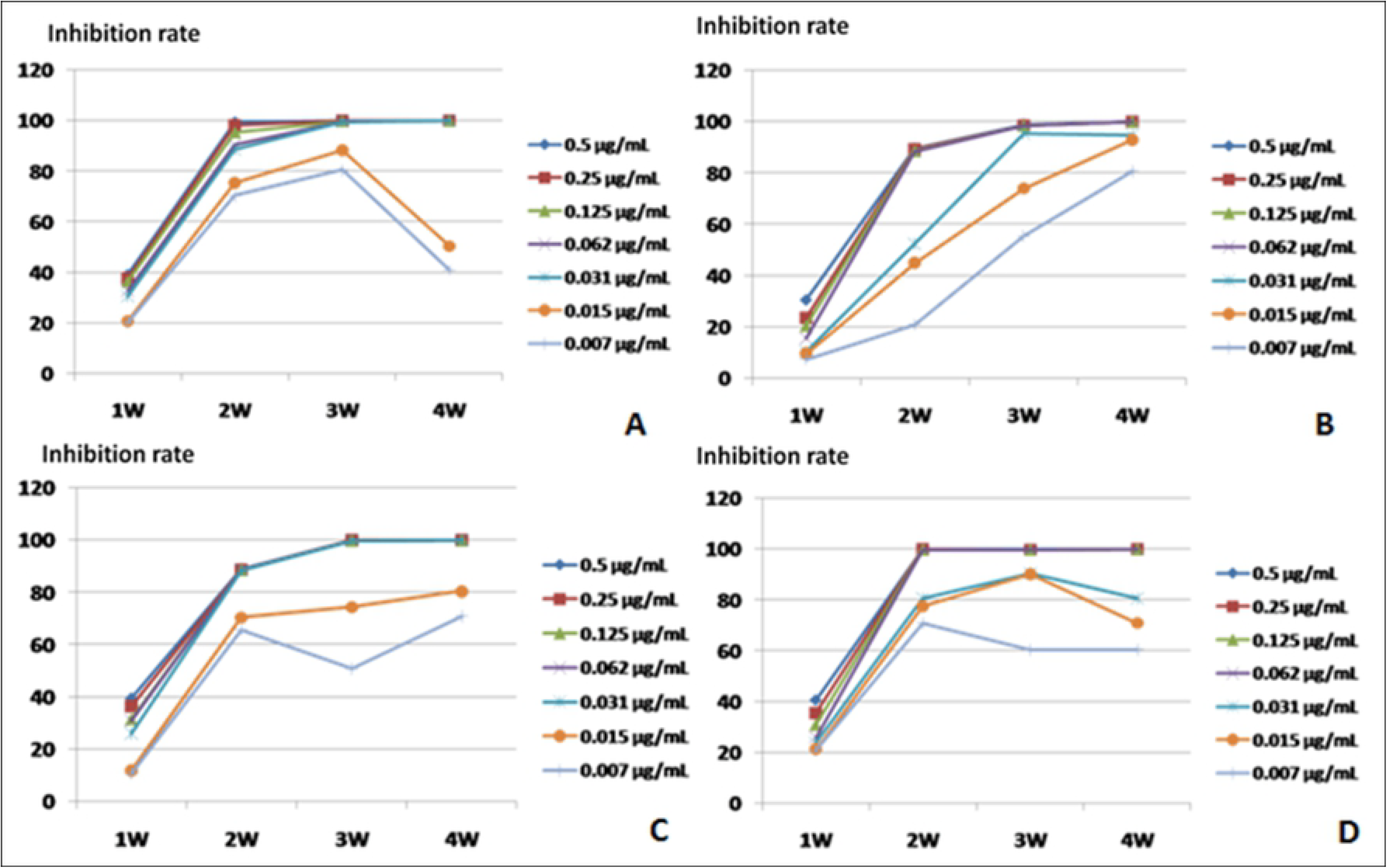
Growth inhibition of MG1 (A), MG2 (B), MG3 (C) and MS (D) grown into mycoplasma medium with various concentrations of tylvalosin at 1, 2, 3 and 4weeks of incubation.

The CT values of mycoplasma isolates in medium inoculated with tested substances increased with increasing the efficacy of such antimicrobials, implying that the mycoplasma loads decreased. A remarkable reduction in the DNA loads of both MG *mgc2* and MS *vlh1* genes was observed following treatment with majority of the screened substances (up to 0.0014 and 0.010, respectively). DNA copy numbers of both MG and MS *mgc2* and *vlh1* gene copies were markedly decreased upon treatment with tylvalosin (up to 0.0032). Therefore, a representative model of growth inhibition rates of MG and MS isolates after exposure to various concentrations of tylvalosin at 1^st^, 2^nd^, 3^th^ and 4^th^ weeks of incubation is shown in figure 1. *Mycoplasma synoviae* isolate had a rapid growth pattern as MIC for tylvalosin was determined early at 2 weeks after incubation. The MIC value was 0.062 µg/mL at 2^nd^, 3^rd^ and 4^th^ weeks of incubation. All MG isolates grew slowly as MICs couldn’t be detected at 1^st^ and 2^nd^ weeks. For MG1 and MG3, the MIC value was 0.031µg/mL at both 3^rd^ and 4^th^ weeks and for MG2, the MIC was 0.062 µg/mL at the 4^th^ week of incubation. Based on this model, MIC values of other examined antimicrobials were determined and shown in Tables 2 and 4.

#### Comparison between conventional broth microdilution and qRT-PCR assays for MICs determination of tested antimicrobials

In Tables 2 and 4, a comparison between all MIC results obtained by conventional broth microdilution and real time PCR methods was carried out. The MICs determined by qRT-PCR assay were similar to the initial MICs by the conventional method for lincomycin (median, 1.5 versus 1.5 µg/mL), enrofloxacin (median, 0.5 versus 0.5 µg/mL), tylosin (median,0.125 versus 0.06 µg/mL), doxycycline (median, 0.25 versus 0.19 µg/mL), tiamulin (median, 0.38 versus 0.19 µg/mL), clove oil (median, 0.74 versus 0.98 µg/mL) and toldin (median, 0.49 versus 0.74 µg/mL), but higher for chlortetracycline (median, 4 versus 1.5 µg/mL), cinnamon (median, 3.91 versus 5.86 µg/mL) and cumin (median, 0.98 versus 1.47 µg/mL).

## Discussion

Avian mycoplasmosis is one of the most serious infectious diseases caused by *Mycoplasma* species, especially MG and MS. Appropriate treatment is recommended in the control of these infections in order to minimize their economic losses. *In vitro* antimicrobial sensitivity testing of avian *Mycoplasma* species has been carried out using broth microdilution method. Meanwhile, we are facing the absence of researches about the inhibitory effects of various antimicrobial substances on clinical MG and MS strains by the use of qRT-PCR as a sensitive method. Therefore, this is the first work that evaluated the *in vitro* susceptibilities of MG and MS isolated from chicken and turkey flocks in Egypt against various antibiotics and essential oils using both conventional broth microdilution method and qRT-PCR.

In the present study, phenotypic and genotypic identification of the recovered isolates revealed high isolation rates of *Mycoplasma* species from the cultured tracheal swabs of the examined chicken (29.3%) and turkey (30%) flocks. Our findings are close to those previously observed in other recent researches in Egypt, where high overall recovery rates of *Mycoplasma* species from chicken tracheal swabs (55.65%) [16] and from diseased turkey flocks (24%) [17] were recorded. Another research conducted inEastern Algeria reported also a high isolation rate of *Mycoplasma* species from tracheas of dead chickens (27%) [18]. The high mycoplasma prevalence rates in these investigated flocks may refer to the already acknowledged vertical transmission characteristics of mycoplasmas in addition to their horizontal transmission to the healthy susceptible birds, which amplifies the disease incidence and the subsequent economic losses. In terms of MG, the isolation rates from the diseased chickens and turkeys were 27.33 and 30%, respectively. In this regard, MG was previously isolated from 40 and 20% of chicken and turkey flocks, respectively [19]. Moreover, in other researches performed earlier, MG prevalence rates from tracheal swabs collected from broilers and turkeys were 38.71% [16] and 68% [20], respectively. With regards to MS, a lower incidence rate was recorded from investigated chicken flocks (2%), although it could not be isolated previously from broilers in Iran [21]. On the other hand, several researchers reported higher prevalence rates of MS from broiler chickens in Iran (31.25%) [22], Jordan (21.7%) [23] and Tehran province (17.89%) [24].

From our findings, cinnamon, cumin, clove oil and Toldin CRD had various antimycoplasmal activities from good to excellent with MIC values up to 0.49 µg/mL. The potent activities observed for plant extracts against mycoplasma organisms may be connected to the anatomical structure of mycoplasma which lacks a rigid cell wall. This feature enables the plant extracts to diffuse into the mycoplasma microorganisms, thereby causing inhibition of growth or death of mycoplasmas.

The extracts were analyzed by the previous established criteria which determines that substances with MIC values < 10 μg/mL are considered to have an excellent antibacterial activity, values between 10 and 100 μg/ mL are considered good, values between 100 and 500 μg/mL are considered moderate, values between 500 and 1000 μg/mL are considered weak and the substances are considered inactive when MIC values are above 1000 μg/mL [8].

Toldin CRD had the best activity with an MIC of 0.49-0.98 µg/mL for all examined isolates. To our best knowledge, the antibacterial effects of Toldin CRD have not been described yet. It is possible that, the strong antimycoplasmal activity of Toldin CRD may be explained by the presence of many active compounds which are powerful bacteriolytic agents such as aliumcepa, ginger, cinnamon, oregano, circumin, liqurice and anise oils and propolis. The antibacterial activities of these compounds were identified previously against a large variety of pathogenic bacteria [8,25].

Strong antimycoplasmal capacities were also determined for both clove and cinnamon oils against both MG and MS isolates. In connection with previous researches performed on other *Mycoplasma* species, euganol; a major ingredient of the clove oil and cinnamon possessed bactericidal activities against *M. hominis* in Czech Republic [26]. One of the most remarkable results of the present study was the pronounced antibacterial property of cumin against screened mycoplasma isolates with an MIC value of 0.98-3.91 µg/mL. There are several studies affirmed that cumin exhibited a significant antimicrobial activity against pathogenic bacteria including *Bacillus subtilis*, *E. coli*, *S. aureus*, *P. aeruginosa*, *K. pneumonea*, *Strept. feacalis* and *S*. Typhimurium [27]. However, this is the first research that evaluates the antibacterial activity of cumin against bacteria without cell wall like mycoplasma.

As our data show (Table 4), conventional microdilution results exhibited a remarkable deviation when initial and final readings were compared. Only in the case of 6 antibiotics (tylvalosin, tilmicosin, doxycycline, chlortetracycline, lincomycin and tiamulin), initial and final MICs exhibited up to 3 fold differences among MG isolates, although it did not alter the interpretation of the data. On the other hand, final MICs for enrofloxacin and lincomycin differed from those of the initial MICs among each of MG2 and MS isolates. The observed MIC differences of these antibiotics lead to the re-categorization of these isolates from susceptible to resistant during interpretation of the results. These results are consistent with a previous report carried out in Germany, where the final MICs for enrofloxacin differed from that of the initial MICs including changes of the profile from sensitive to resistant among most of MS isolates [4]. The discrepancies may be attributed to the inactivation of the used antibiotics with the continued incubation time [28] or the presence of a mixed colony in the field isolates with a slower growing minor population [13,29]. Therefore, the initial MIC values are advised to be taken into consideration in the interpretation of the susceptibility results due to these values compare the growth of each mycoplasma isolate in the presence and absence of tested substances rather than determining the endpoint at a fixed time [13].

According to our results, various susceptibility rates of MG and MS isolates were observed against the eight antibiotics. Macrolides proved to have good *in vitro* effectiveness against both MG and MS isolates with lower MIC values ranging from 0.015-0.25 µg/mL. Lower MIC values for macrolides were also reported in previous examinations [5,17] supporting the use of these antibiotics for treatment of mycoplasmas infecting poultry in the field.

Among macrolides group, tylvalosin was the most effective drug against both MG and MS isolates with an MIC range of 0.015-0.030 µg/mL. This finding is in a complete agreement with other research results in Iran [30], UK [5] and Europe [15]. This could be attributed to the recent introduction of tylvalosin as a newer antibiotic against MG and MS isolates in Egypt. Our field experience supports this finding.

In the present investigation, DNA copies of both MG *mgc2* and MS *vlhA* genes were markedly decreased upon treatment with majority of the tested antimicrobials with relevant MIC values determined by qRT-PCR assay confirming the good effectiveness of these compounds. The MICs determined using qRT-PCR were almost similar to those obtained by conventional microdilution method. Only, a previous report in Japan was carried out to determine antibiotic susceptibility testing of *M. genitalium* by TaqMan 5’ nuclease qRT-PCR assay [14]. Recently, assessing the inhibitory effects of antibiotics using qRT-PCR assay has been also described on common clinically relevant bacterial species such as *S. aureus*, *E. coli* and *Haemophilusinfluenzae* in France [9].

## Conclusion

The current study showed that Toldin CRD and tylvalosin possessed strong *in vitro* antimycoplasmal activities against both MG and MS field isolates. The MIC values determined using qRT-PCR were similar to those obtained by the conventional microdilution method. However, molecular detection of antimicrobial susceptibility using qRT-PCR could be more sensitive than phenotypic methods.

